# The Defensive Activation theory: dreaming as a mechanism to prevent takeover of the visual cortex

**DOI:** 10.1101/2020.07.24.219089

**Authors:** David M. Eagleman, Don A. Vaughn

## Abstract

Regions of the brain maintain their territory with continuous activity: if activity slows or stops (e.g., because of blindness), the territory tends to be taken over by its neighbors. A surprise in recent years has been the speed of takeover, which is measurable within an hour. These findings lead us to a new hypothesis on the origin of dream sleep. We hypothesize that the circuitry underlying dreaming serves to amplify the visual system’s activity periodically throughout the night, allowing it to defend its territory against takeover from other senses. We find that measures of plasticity across 25 species of primates correlate positively with the proportion of rapid eye movement (REM) sleep. We further find that plasticity and REM sleep increase in lockstep with evolutionary recency to humans. Finally, our hypothesis is consistent with the decrease in REM sleep and parallel decrease in neuroplasticity with aging.

## Introduction

One of neuroscience’s unsolved mysteries is why brains dream (*1*–*4*). Do our bizarre nighttime hallucinations carry meaning, or are they simply random neural activity in search of a coherent narrative? And why are dreams so richly visual, activating the occipital cortex so strongly? We here leverage recent findings on neural plasticity to propose a novel hypothesis.

Just as sharp teeth and fast legs are useful for survival, so is neural plasticity: the brain’s ability to adjust its parameters (e.g., the strength of synaptic connections) enables learning, memory, and behavioral flexibility (*5*–*7*).

On the scale of brain regions, neuroplasticity allows areas associated with different sensory modalities to gain or lose neural territory when inputs slow, stop, or shift. For example, in the congenitally blind, the occipital cortex is taken over by other senses such as audition and somatosensation (*8*). Similarly, when human adults who recently lost their sight listen to sounds while undergoing functional magnetic resonance imaging (fMRI), the auditory stimulation causes activity not only in the auditory cortex, but also in the occipital cortex (*9*). Such findings illustrate that the brain undergoes changes rapidly when visual input stops.

Rapid neural reorganization happens not only in the newly blind, but also among sighted participants with temporary blindness. In one study, sighted participants were blindfolded for five days and put through an intensive Braille-training paradigm (*10*). At the end of five days, the participants could distinguish subtle differences between Braille characters much better than a control group of sighted participants who received the same training without a blindfold. The difference in neural activity was especially striking: in response to touch and sound, blindfolded participants showed activation in the occipital cortex as well as in the somatosensory cortex and auditory cortex, respectively. When the new occipital lobe activity was intentionally disrupted by magnetic pulses, the Braille-reading advantage of the blindfolded subjects went away. This finding indicates that the recruitment of this brain area was not an accidental side effect—it was critical for the improved performance. After the blindfold was removed, the response of the occipital cortex to touch and sound disappeared within a day.

Of particular interest here is the unprecedented speed of the changes. When sighted participants were asked to perform a touching task that required fine discrimination, investigators detected touch-related activity emerging in the primary visual cortex after only 40 to 60 minutes of blindfolding (*11*). The rapidity of the change may be explained not by the growth of new axons, but by the unmasking of pre-existing non-visual connections in the occipital cortex.

It is advantageous to redistribute neural territory when a sense is permanently lost, but the rapid conquest of territory may be disadvantageous when input to a sense is diminished only temporarily, as in the blindfold experiment. This consideration leads us to propose a new hypothesis for the brain’s activity at night. In the ceaseless competition for brain territory, the visual system in particular has a unique problem: due to the planet’s rotation, we are cast into darkness for an average of 12 hours every cycle. (This of course refers to the vast majority of evolutionary time, not to our present electrified world). Given that sensory deprivation triggers takeover by neighboring territories (*12*–*14*), how does the visual system compensate for its cyclical loss of input?

We suggest that the brain combats neuroplastic incursions into the visual system by keeping the occipital cortex active at night. We term this the Defensive Activation theory. In this view, dream sleep exists to keep the visual cortex from being taken over by neighboring cortical areas. After all, the rotation of the planet does not diminish touch, hearing, taste, or smell. Only visual input is occluded by darkness.

In humans, sleep is punctuated by REM (rapid eye movement) sleep about every 90 minutes (*15*). This is when most dreaming occurs. Although some forms of dreaming can occur during non-REM sleep, such dreams are quite different from REM dreams; non-REM dreams usually are related to plans or thoughts, and they lack the visual vividness and hallucinatory and delusory components of REM dreams (*16*).

REM sleep is triggered by a specialized set of neurons in the pons (*17*). Increased activity in this neuronal population has two consequences. First, elaborate neural circuitry keeps the body immobile during REM sleep by paralyzing major muscle groups (*18*). The muscle shut-down allows the brain to simulate a visual experience without moving the body at the same time. Second, we experience vision when waves of activity travel from the pons to the lateral geniculate nucleus and then to the occipital cortex (these are known as ponto-geniculo-occipital waves or PGO waves) (*19*). When the spikes of activity arrive at the occipital pole, we feel as though we are seeing even though our eyes are closed (*3*). The visual cortical activity is presumably why dreams are pictorial and filmic instead of conceptual or abstract.

These nighttime volleys of activity are anatomically precise. The pontine circuitry connects specifically to the lateral geniculate nucleus, which passes the activity on to the occipital cortex, only. The high specificity of this circuitry supports the biological importance of dream sleep: putatively, this circuitry would be unlikely to evolve without an important function behind it.

The Defensive Activation theory makes a strong prediction: the higher an organism’s neural plasticity, the higher its ratio of REM to non-REM sleep. This relationship should be observable across species as well as within a given species across the lifespan. We thus set out to test our hypothesis by comparing 25 species of primates on behavioral measures of plasticity and the fraction of sleep time they spend in REM.

## Methods

We first collected published measures of REM sleep (specifically, the fraction of sleep time spent in REM) for 25 species of primate (**Table 1 and Supplemental Material**). All data was collected on juvenile and adult primates; no data from infants or very young primates was available. We resolved discrepancies between publications by averaging the values.

**Table 1.** Three behavioral correlates of plasticity and the proportion of REM (versus non-REM) sleep in 25 primate species. All references in Supplemental Material.

| Coloquial Name | Time to Locomotion (Days) | Time to Weaning (Days) | Time to Adolescence (Months) | Percentage of Sleep in REM | Phylogenetic Distance from Humans (M yrs) |
| --- | --- | --- | --- | --- | --- |
| vervet monkey | 21 | 117 | 48 | 6% | 29 |
| mongoose lemur | 35 | 150 | 36 | 7% | 74 |
| grey mouse lemur | 21 | 51 | 2 | 7% | 74 |
| spider monkey | — | 821 | — | 7% | 43 |
| yellow baboon | 60 | 365 | 16 | 8% | 29 |
| patas monkey | 150 | 216 | 22 | 9% | 29 |
| black lemur | 43 | 170 | 18 | 9% | 74 |
| barbary macaque | — | 258 | 12 | 9% | 29 |
| senegal galago | 7 | 105 | 2 | 9% | 74 |
| green monkey | — | 365 | 12 | 9% | 29 |
| brown lemur | — | 150 | 25 | 9% | 74 |
| guinea baboon | 14 | 420 | 14 | 11% | 29 |
| three-striped night monkey | 34 | 77 | 42 | 11% | 43 |
| long-tailed macaque | — | 385 | — | 11% | 29 |
| pigtail macaque | 35 | 342 | 11 | 12% | 29 |
| bornean orangutan | 120 | 1637 | 60 | 12% | 16 |
| hamadryas baboon | — | 315 | 24 | 13% | 29 |
| olive baboon | 14 | 390 | 14 | 13% | 29 |
| bonnet macaque | 180 | 195 | 11 | 14% | 29 |
| common marmoset | 14 | 81 | 5 | 14% | 43 |
| stump-tailed macaque | — | 270 | 18 | 15% | 29 |
| common squirrel monkey | 160 | 177 | 24 | 15% | 43 |
| chimpanzee | 70 | 1393 | 96 | 16% | 7 |
| rhesus monkey | 180 | 365 | 78 | 18% | 29 |
| human | 318 | 1095 | 162 | 21% | 0 |

Next, we collected measures of plasticity for the same species. Ideally, we would analyze the plasticity of biochemical and genetic changes in the visual cortex, such as long term potentiation or epigenetic modifications. However, because such data are extremely sparse across species, we instead employed three behavioral proxies of plasticity across 25 primates: time to weaning, time to locomotion, and time to adolescence (**Table 1**). Most data on time to weaning and time to adolescence were collected from the Human Ageing Genomic Resources website (genomics.senescence.info), which aggregates data from published studies, and we supplemented as needed with Animal Diversity Web (animaldiversity.org) to find values for all 25 species. Locomotion data were taken from Primate Info Net by the University of Wisconsin (pin.primate.wisc.edu) and Animal Diversity Web, though we were able to locate values for only 18 of the 25 primates. Phylogenetic distances were gathered from timetree.org, which aggregates data from published studies.

## Statistical Methods

Significance values were obtained using an F-test of a model with plasticity variable(s) versus a model with only a constant. We corrected for multiple comparisons on multiple univariate correlations using the Holm-Bonferroni method.

We detected distributional non-normality using the Shapiro-Wilk test, and we computed the Shapiro-Wilk statistic for both the initial plasticity variables and the residuals from each regression. The null hypothesis (that the data are normally distributed) is retained unless otherwise stated.

The plasticity metric distributions were skewed rightward and non-normal (time to locomotion, W = 0.80, *p* < 0.001; time to weaning, W = 0.71, *p* < 10^−4^; and time to adolescence, W = 0.71, *p* < 10^−4^). To prevent the undue influence of outliers on univariate correlation values, we used a logarithmic transformation (base 10) of all three plasticity metrics.

We computed a plasticity index by performing principal component analysis (PCA) on the three behavioral expressions of plasticity. We employed an ordinary least squares multivariate regression on the three behavioral metrics of plasticity to predict REM values. Given the results of the Shapiro Wilk test above, all three features were transformed into log space for both the PCA and the regression.

## Results

Does the general plasticity of a species correlate with the percentage of sleep time they spend in REM? We found a significant correlation with all three measures (corrected for multiple comparisons): time to locomotion (**Fig 2A**, *r*^2^ = 0.32, *F*(17) = 7.4, corrected *p* < 0.05), time to weaning (**Fig 2B**, *r*^2^ = 0.17, *F*(24) = 4.8, corrected *p* < 0.05), and time to adolescence (**Fig 2C**, *r*^2^ = 0.22, *F*(22) = 5.9, corrected *p* < 0.05). A single multivariate model incorporating all three behavioral proxies of plasticity performed significantly better than chance (**Fig 2D**, *r*^2^ adjusted = 0.29, *F*(14) = 3.3, *p* < 0.05).

**Fig. 1.**
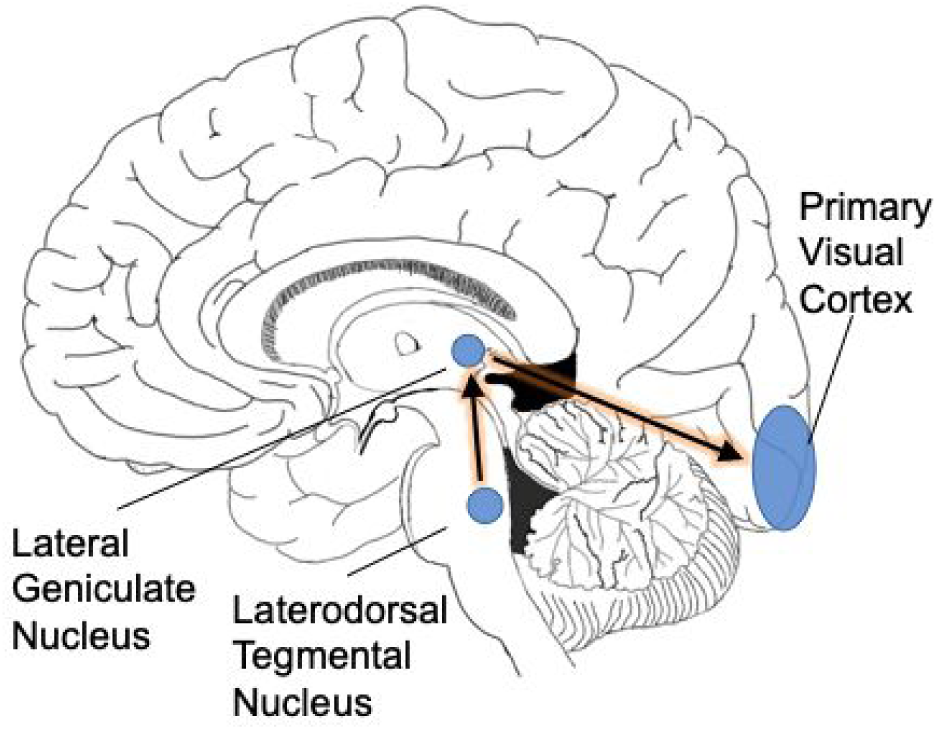
PGO waves. As a prelude to REM sleep, waves of activity move from the brainstem into the occipital cortex. We suggest that this infusion of activity is necessitated by the rotation of the planet into darkness: the visual system needs extra cyclic activation to keep its territory intact.

**Fig. 2.**
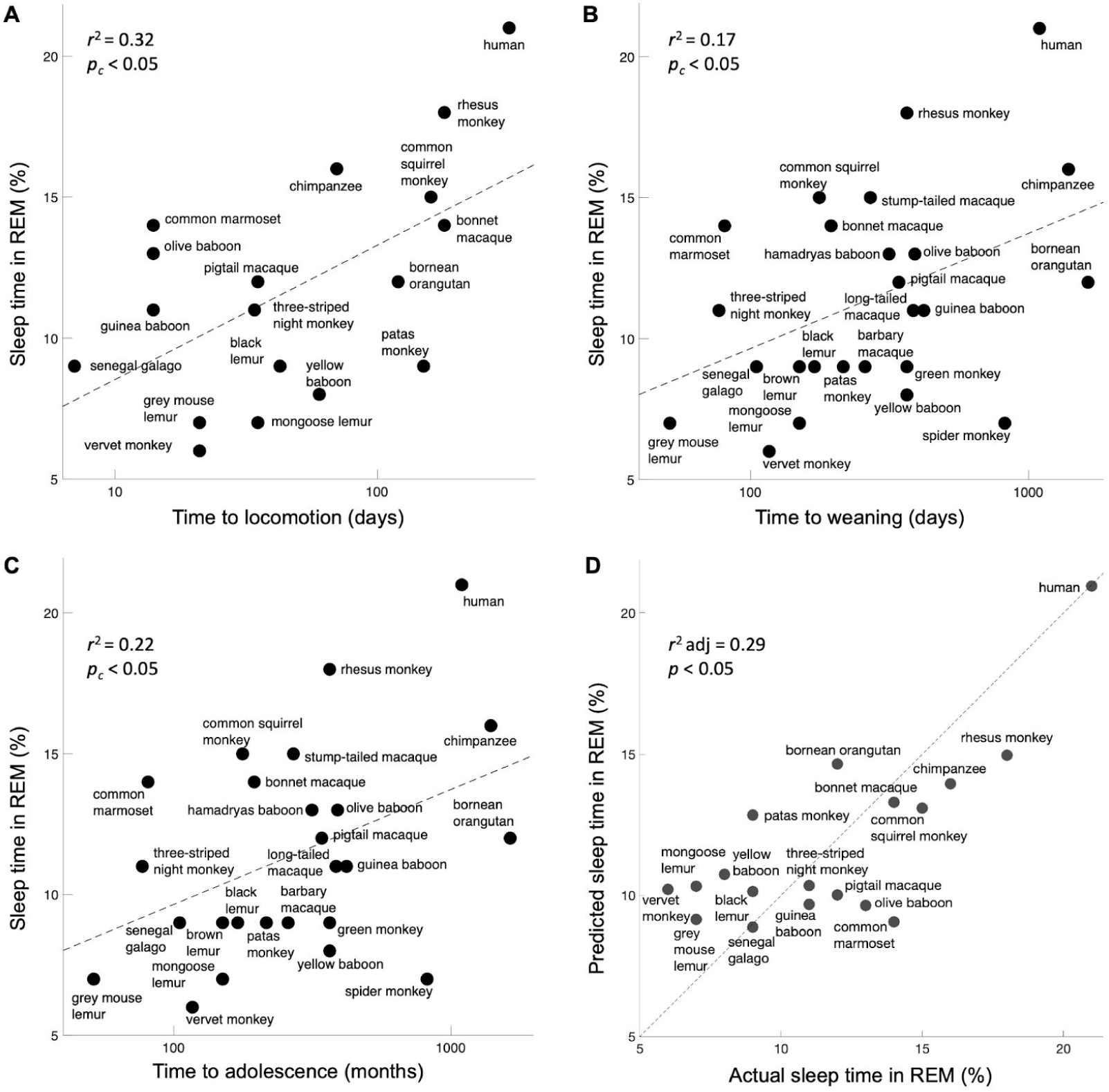
Expressions of plasticity correlate with the percentage of sleep time spent in REM. **(A)** Time to locomotion. **(B)** Time to weaning. **(C)** Time to adolescence. Although these are crude correlates of neuroplasticity, they map significantly onto the fraction of dream sleep, as per the prediction. **(D)** A model incorporating all three behavioral measures of plasticity significantly predicts the percentage of sleep time spent in REM. The unity line signifies the perfect model prediction.

Our data show that more plasticity correlates with more REM sleep. These two measures are putatively connected by the necessity of defending neural territory that cyclically lacks external stimulation. As a next step, we plotted the same measures of plasticity and REM sleep as a function of phylogenetic distance from humans (**Figure 3A**). The data demonstrate that both plasticity and REM sleep correlate significantly with the phylogenetic distance from humans (corrected *p* < 0.001 and *p* < 0.01, respectively). This finding suggests that plasticity increases as brain complexity increases, perhaps because higher-dimensional systems require more training data to converge (e.g., in machine learning (*20*)) or because greater complexity otherwise would require a numerical explosion of genes to control the operation of intricate cortices. With that result in mind, we next assessed whether complexity, operationalized as the number of cone inputs, varies with REM sleep. Indeed, monochromatic and dichromatic primates, who split off from the lineage of *Homo sapiens* many tens of millions of years ago, spent a smaller proportion of sleep time in REM (**Figure 3B**, *p* < 0.05, *t*(23) = 2.2, two-tailed t-test).

**Fig. 3.**
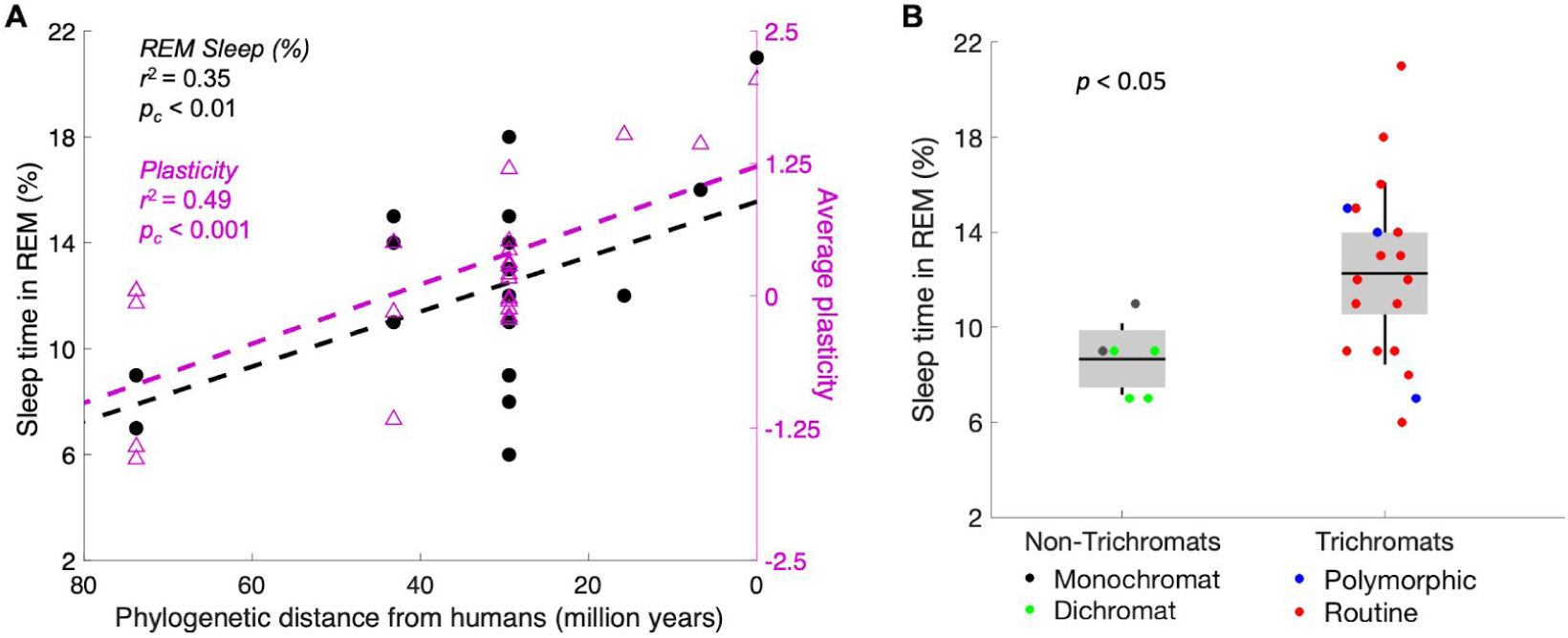
Plasticity and REM sleep increase concomitantly with evolutionary recency to humans. **(A)** Primates are plotted by recency of common ancestors from humans, with percentage of sleep time in REM (left axis) and behavioral plasticity index (right axis). The correlation is significant for both measures. **(B)** Trichromats average 49% more REM sleep (as a fraction of total sleep time) than monochromats and dichromats. We hypothesize that the increased complexity of the visual system correlates with higher plasticity, which then requires more defense of the visual system during the night.

Within a given species, our hypothesis predicts that REM sleep decreases across the lifespan because neuroplasticity decreases with age. For example, infants’ brains are more plastic, and thus the competition for territory is even more critical. As an animal ages, the decrease in neuroplasticity makes cortical takeover increasingly difficult or impossible (*21*), so the proportion of sleep time spent in REM should decrease with age. The data are consistent with this prediction: there is a clear and significant decrease in REM (as a percentage of total sleep time) across the lifespan (*15*). In humans, for example, REM accounts for half of infants’ sleep time but only 10-20% of adults’ sleep time; the percentage decreases slowly from young adults to elderly adults (**Figure 4A**). Note that this trend is directionally consistent with the decrease in plasticity during the human lifespan, as measured by studies on memory skills (*22*), motor cortex plasticity (*23*) (**Figure 4B**), calcium conductance in neurons (*24*), synaptic response size (*25*), long-term potentiation (*26*), and many more (*27*). We suggest that the directionality is not coincidental—rather, REM sleep becomes less necessary as neural circuitry becomes less flexible.

**Fig. 4.**
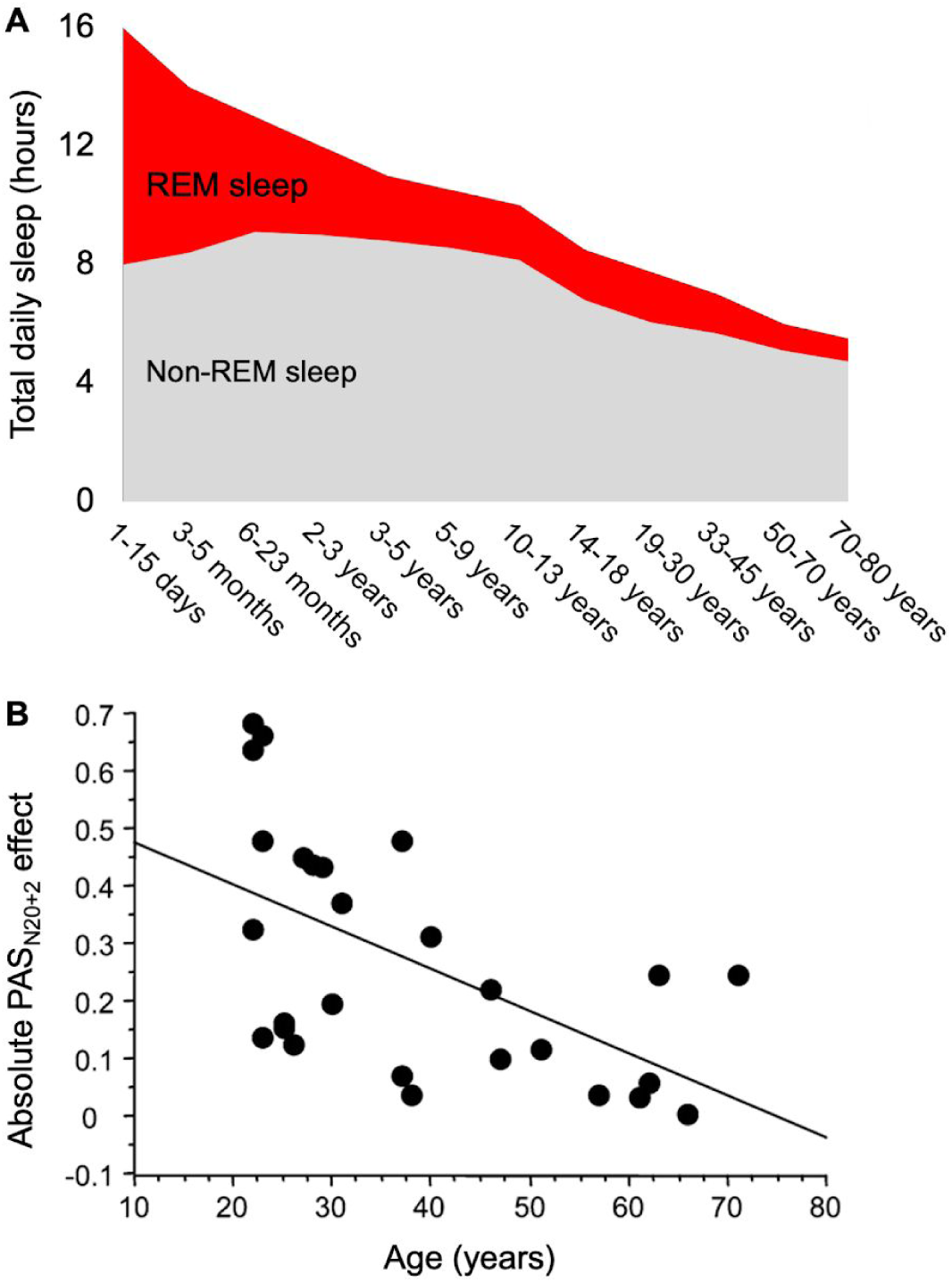
The decrease in REM sleep with age parallels the decrease in plasticity. This result is directionally consistent with the present hypothesis that REM sleep is less necessary when neural circuitry is less flexible. **(A)** Across animal species, the fraction of sleep time spent in REM decreases (data in humans adapted from (28)). **(B)** As just one example (see text for others), neuroplasticity diminishes with age in the human motor cortex, here measured with a paired associative stimulation protocol. Data from (23).

The decline in plasticity is consistent with the aging brain’s declining ability to recover from damage. For example, the amount of takeover in the occipital cortex is greatest if a person is born blind; takeover is modest if blindness occurs in youth and is less pronounced if blindness occurs later in life (*29*). The same trend is seen with damage sustained from strokes: the younger the person, the better the chance of recovery via plasticity (*30*).

## Discussion

We have proposed a new hypothesis about the purpose of REM sleep: specifically, that it may defend the visual system against cortical takeover from other senses during sleep (**Figure 5**). To evaluate a potential correlation between REM sleep and plasticity, we compared three measures of plasticity across 25 primate species, including humans. Results (**Figure 2**) demonstrate the plausibility of our hypothesis as all three behavioral expressions of plasticity correlate significantly with the fraction of sleep time spent in REM.

**Fig. 5.**
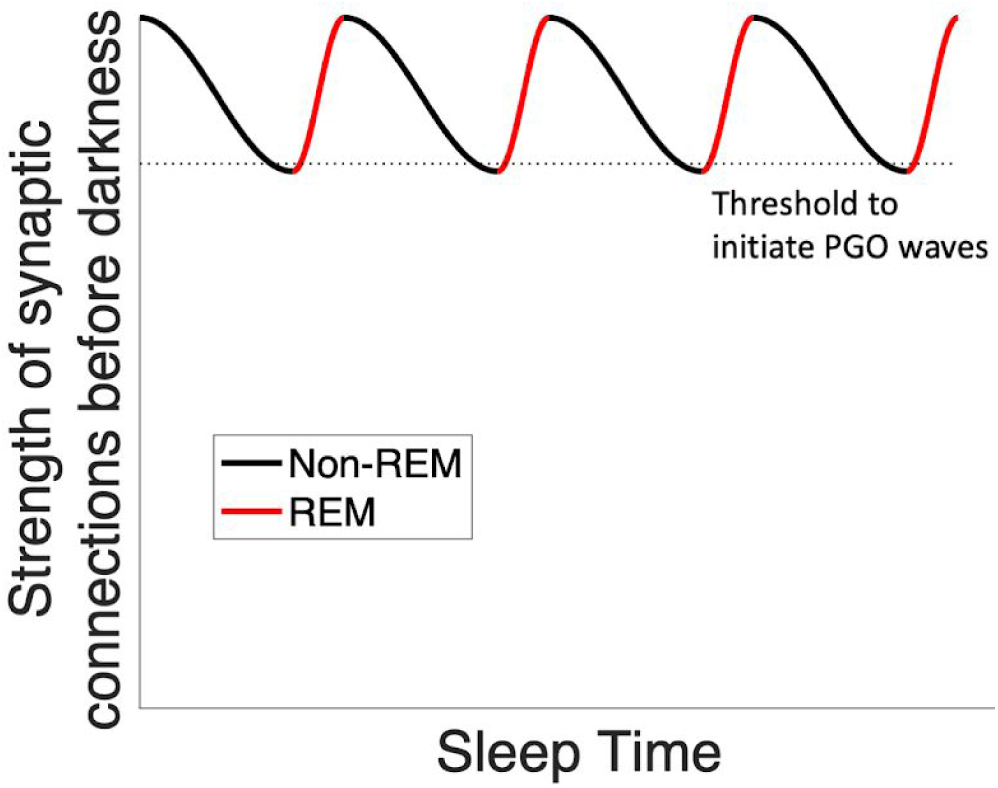
Representation of the Defensive Activation theory. With the onset of sleep, visual synaptic connections are weakened by encroachment from other sensory areas. When a threshold is reached (dotted line), PGO waves are initiated and drive activity into the occipital lobe. This process repeats cyclically throughout the sleep cycle.

Eight percent (2/25) of the primate species in our dataset were nocturnal. Note that this does not change the hypothesis: whenever an animal sleeps, whether at night or during the day, the occipital cortex is at risk of takeover by the other senses. Nocturnal primates, equipped with strong night vision, employ their visual cortex throughout the night as they seek food and avoid predation. When they subsequently sleep during the day, their closed eyes allow no visual input, and thus their occipital cortex requires defense.

We further noted that the fraction of REM sleep increases with phylogenetic complexity (**Figure 3**), suggesting the possibility that increasing complexity requires more plasticity (e.g., as opposed to genetic predetermination). Thus, our hypothesis may be a specific example of a general principle: more plastic systems require more active maintenance.

Finally, we noted that as plasticity decreases across the lifespan, so too does the fraction of REM sleep (**Figure 4**). This occurs both in humans and across the animal kingdom (*31*).

It is important to recognize that the present hypothesis must be considered in the context of evolutionary time. Because the circuitry for dreaming presumably evolved over tens of millions of years, it is unaffected by our modern ability to defy darkness with electrical light. It thus comes as no surprise that people born blind retain the same PGO circuitry as the sighted (*32*), and they dream. However, the congenitally- and early-blind experience no visual imagery in their dreams but do have *other* sensory experiences, such as feeling their way around a rearranged living room or hearing strange dogs barking (*33, 34*). This is consistent with the fact that the other senses take over the occipital cortex of the blind (*12, 35*). Thus, in a congenitally blind person, nighttime occipital activation still occurs, but the established circuitry is non-visual. Note also that people who become blind *after* the age of seven have more visual content in their dreams than those who become blind at younger ages (*33*). This is consistent with the finding that the occipital lobe in the late-blind is less fully conquered by the other senses (*9*).

### Methods to test the hypothesis

The present hypothesis could be tested more thoroughly with direct measures of cortical plasticity. An ideal experiment would be to blindfold awake animals and measure takeover of the visual cortex by other senses in functional imaging (e.g., fMRI).

We predict that less-plastic species (e.g., the Senegal galago, from whom humans diverged about 74 million years ago) would show a slower and smaller takeover when compared to humans, in which auditory and tactile cues cause activation in the occipital cortex within about an hour of blindfolding (*11*). Unfortunately, this type of data has been collected only in humans.

Beyond direct imaging, other measures of plasticity could add additional clarification and correlational support. Potential measures include paired associative stimulation or long-term potentiation. Again, at present, such data is sparse across different primate species. Ideally, this line of research would carefully track the developmental stage (e.g., the percentage of sleep time in REM differs between adolescents and adults, a point not tracked consistently in the extant literature.)

Despite the limitations of the present data, our hypothesis can nonetheless be tested more rigorously in humans by examining visual changes in diseases that lead to the loss or impairment of dreaming. For example, REM sleep can be suppressed by monoamine oxidase inhibitors (MAOIs) (*36*) and by a variety of brain lesions (*37*). It remains contentious whether general cognitive or physiological problems occur in people with deficits in REM sleep (*38*), but our hypothesis specifically predicts visual problems, and this is precisely what is seen in patients on MAOIs or tricyclic antidepressants (TCAs). Patients on these medications characteristically experience blurry vision (*39*). This is typically marked up to dry eyes, but we here note our alternative (or complimentary) hypothesis.

Given these possibilities, we suggest two tests for future research. First, one could perform neuroimaging (fMRI) across time with two groups: (1) participants on REM-reducing medications (such as MAOIs or TCAs) and (2) control participants with normal REM sleep. Our hypothesis predicts that the medicated group would show a cyclical change in the activity of the occipital cortex; there would be less visual cortical activity after waking up than before going to sleep. This effect might also be seen in the control group, but we would predict a smaller cyclical change than in the medicated group.

Second, REM sleep could be disrupted non-medically. Volunteers could undergo contingent awakening (in a dark room) followed by imaging of the strength of activity of other senses (e.g., touch and hearing) in the occipital cortex.

### Other species

A logical next step in evaluating this hypothesis would be a systematic comparison of our measures across a variety of animal species (i.e., beyond primates). Some mammals are born immature (altricial), unable to regulate their own temperature, acquire food, or defend themselves. Examples include opossums, ferrets, and platypuses. Other mammals such as guinea pigs, sheep, and giraffes are born mature (precocial), emerging from the womb with teeth, fur, open eyes, and the abilities to regulate their temperature, walk within an hour of birth, and eat solid food. Altricial animals have a greater proportion of REM sleep than precocial animals—up to about 8 times as much—and this difference is especially pronounced in the first month of life (*40, 41*). In the framing of this hypothesis, when a neonatal brain has high plasticity, the system requires more effort to defend the visual system during sleep. Conversely, when a newborn’s brain arrives mostly organized (and less plastic), there is less need to fight encroachment as the system is more immutable.

The study of sleep duration across a wide variety of species is complexified by the fact that sleep duration is modified by many environmental factors, including predation, weather, and sleep location (*42*–*45*). For example, terrestrial mammals have REM sleep but aquatic mammals do not, presumably because their environments require them to move continuously; thus, evolution seems to have de-selected REM sleep in these mammals (*46*). It is still possible that aquatic mammals activate their visual cortex during sleep, but it does not seem to occur in a way that is easy to measure (e.g., rapid eye movement). It is also possible that unihemispheric sleep provides enough cross-hemisphere activation that neither visual cortex is disadvantaged, obviating the need for dreaming. Whatever the case, evolutionary pressures provide many pushes and pulls for solutions, so the story is likely to be confounded with other factors as we look across the animal kingdom.

As another example, consider the surprisingly small amount of REM sleep—a few minutes at most—in the elephant (*47, 48*). At first blush, this seems like a refutation of the present hypothesis. However, note that elephants sleep very little, around two hours a night, and they have excellent night vision due to specializations in their retinas (*49*). As a result, their visual cortex, active during almost all hours of the day and night, does not face the same threat of encroachment from the other senses. Thus, our hypothesis predicts that elephants should have little to no REM sleep. Additionally, we suggest a possible prediction about when elephants do and do not display REM sleep: the specialized retina still requires some light, so elephants suffer poor night vision in the absence of moonlight (*49*). Thus, if REM sleep is partially responsive to outside conditions, our framework predicts more REM sleep in elephants on nights with heavy cloud cover or a new moon as compared to on clear nights with a full moon. It should be noted that a positive result would buttress our hypothesis; the absence of a correlation would simply suggest that the amount of REM sleep may not be responsive to immediate environmental conditions; and an opposite result would stand against our hypothesis.

In any case, the current framework may offer insight into the vastly different sleep times among animals, even those within the same phylogenetic order. For example, the golden mantled ground squirrel (*Spermophilis lateralis*) sleeps 15.9 hours a day, while its close cousin, the degu (*Octodott degu*) sleeps only 7.7 hours (*50*). Of note, however, they spend similar fractions of sleep time in REM: 3.0 hours (18.9%) for the former and 0.9 hours (11.7%) for the latter (*50*). Such a similarity is expected for two animals with similar levels of plasticity. In other words, while many factors influence the total time spent sleeping, the proportion of time during which the visual cortex must defend itself with REM sleep should remain approximately the same for animals with similar neural complexity. Further research will be required to understand the strength of the present hypothesis across animal species, including species that are nocturnal or crepuscular.

### Cortical maintenance

If dreams are visual hallucinations that prevent neuroplastic encroachment, we might expect to find similar visual hallucinations while awake, during prolonged periods of sensory deprivation. In fact, visual hallucinations have been reported by individuals deprived of novel visual input, such as people with macular degeneration, people confined to a tank-respirator (such as patients with poliomyelitis), patients bedridden after a medical procedure, and prisoners in solitary confinement (*51*). In several of these cases, individuals reported that hallucinations worsened during the nighttime when novel sensory input was further truncated (*52*).

Such phenomena may represent the same mechanism of cortical maintenance. If so, they might give additional insight into the neural architecture of PGO waves. Just as a home thermostat runs not based on the time of day but rather on the ambient temperature of the house it is trying to maintain, PGO waves may be triggered not by circadian rhythms per se but rather by feedback that the visual cortex is experiencing a decline in externally-driven activity. Thus, our anatomical prediction is that axons project from the visual cortex back to the pons, directly or indirectly. If true, then PGO waves would operate as a feedback loop rather than as a one-way directive from the pons to the occipital lobe; they might be better described as OPGO (occipito-pontine-geniculo-occipital) waves. Such a threshold-monitoring circuit would produce cyclical activation of the visual cortex, explaining the cyclical nature of REM (**Figure 5**).

Our Defensive Activation theory—that visual hallucinations serve to counteract neuroplastic encroachment during extended periods of darkness—may represent a more general principle: the brain has evolved specific neural architectures to elicit activity during periods of sensory deprivation. This might occur in several scenarios: when deprivation is regular and predictable (e.g., dreams during sleep), when there is damage to the sensory input pathway (e.g., tinnitus or phantom limb syndrome), and when deprivation is unpredictable and environmental (e.g., hallucinations induced by sensory deprivation).

This hypothesis leads to some testable speculations. In its strongest form, it predicts the possibility of other sensory neural pathways analogous to the PGO pathway. In the case of audition, for example, we suggest there may be PGT (pontine-geniculo-temporal) waves passing through the medial geniculate nucleus rather than the lateral. Such a pathway would have nothing to do with sleep but instead would exist in the (presumably rare) event that input to the auditory cortex slows or stops. However, as deprivations in other senses are ecologically rare, such specialized circuitry may have been evolutionarily unnecessary. In that case, a weaker form of the hypothesis would predict that the local architecture of sensory cortices is structured with feedback loops that manifest random activity when input diminishes. Thus, the neural activity underlying low-level sensory hallucination during periods of deprivation may in fact be a feature of the system rather than a bug.

We have suggested that dream sleep exists at least in part to stave off the takeover of visual territory by other senses during the darkness of night. We propose that this is a necessary maintenance mechanism for neuroplastic systems, and we show that animals with greater plasticity correspondingly spend a greater fraction of sleep time in REM. We tentatively suggest that visual dreaming may be a specific manifestation of a more general principle: that plastic systems have evolved feedback mechanisms to foment activity during periods of sensory deprivation, thus preventing takeover by neural neighbors.

## Supplemental Materials

We collected published measures of REM sleep (specifically, the fraction of sleep time spent in REM) for 25 species of primate, all of whom such measures were available: mongoose lemur (*53, 54*), grey mouse lemur (*53, 55, 56*), vervet monkey (*53, 56, 57*), patas monkey (*53, 57, 58*), black lemur (*54, 56*), barbary macaque (*53, 56, 59*), pigtail macaque (*60*–*62*), guinea baboon (*63*–*65*), three-striped night monkey (*53, 58, 66*), bonnet macaque (*67*–*69*), bornean orangutan (*45*), hamadryas baboon (*56, 70*), olive baboon (*53, 71, 72*), chimpanzee (*58, 73, 74*), stump-tailed macaque (*53, 56, 75*), long-tailed macaque (*76*), common marmoset (*53, 76, 77*), common squirrel monkey (*78*–*80*), rhesus monkey (*65, 81, 82*), human (*83*–*85*), senegal galago (*56, 58, 86*), yellow baboon (*56, 87*), green monkey (*57, 88*), spider monkey (*89*), and brown lemur (*54*).

